# Fructose vs. Glucose: Modulating Stem Cell Growth and Function Through Sugar Supplementation

**DOI:** 10.1101/2024.01.18.576243

**Authors:** Salaheldeen Elsaid, Xiangdong Wu, Sui Seng Tee

## Abstract

In multicellular organisms, stem cells are impacted by microenvironmental resources such as nutrient availability and oxygen tension for their survival, growth, and differentiation^1,2^. However, the acessibility of these resources in the pericellular environment greatly varies from organ to organ^3–5^. This divergence in resource availability leads to variations in the potency and differentiation potential of stem cells^4,6^. Moreover, hexose and oxygen levels modulate cytokine production which is crucial for cell-cell communication^7,8^ as well as growth, and differentiation of stem cell ^9,10^.

Hence, this study aims to explore the distinct effects of glucose and fructose, as well as different oxygen tensions, on the growth dynamics, cytokine production, and differentiation of stem cells.

## Introduction

The availability of sugars and oxygen in the stem cell microenvironment plays a pivotal role in their metabolism, proliferation, and differentiation^1,2^. Notably, these resources are not infinite, but their usage is tightly regulated via cytokines and hormones, that orchestrate a network of differential metabolic fates^9,11^. This includes the activation of membrane transporters and anabolic enzymes, facilitating the uptake of sugars and its conversion into cellular biomass^10,12^. Hence, sugars and oxygen are essential components of cellular metabolism and are crucial for stem cell growth, differentiation, and cytokine expression.

At the same time, abnormally high levels of glucose and fructose in diabetic patients create a substantial stress to stem cells. This can lead to adverse effects on different stem cells by prompting their differentiation into new blood capillaries (angiogenesis) in the retina ^10,12–14^ or adipocytes in bone marrow (adipogenesis)^1^. Therefore, the maintenance of hexoses at physiological levels supports regular metabolic and cellular functions, while its disruption induces a significant stress on stem cells.

Many studies have focused on differentiating stem cells into particular cell types, such as adipocytes^15^, chondrocytes^1,16^, hepatocytes^2^, and osteoblasts^17^. Other studies focused on their advantageous immunomodulatory properties as they secrete intrinsic cytokines that have the potential to mitigate inflammations and attenuate fibrosis^18,19^. Consequently, stem cells are considered indispensable due to their unique combination of multipotency and immunomodulatory capabilities.

Stem cells can be isolated from various tissues that differ not only in their origin but also in their microenvironment. This microenvironment may differ in hexose concentrations and oxygen level^2,16,20^. Consequently, we aim to study the differential effect of different concentrations of glucose and fructose as well as distinct oxygen tensions on stem cells growth dynamics, cytokine production and differentiation.

## MATERIALS AND METHODS

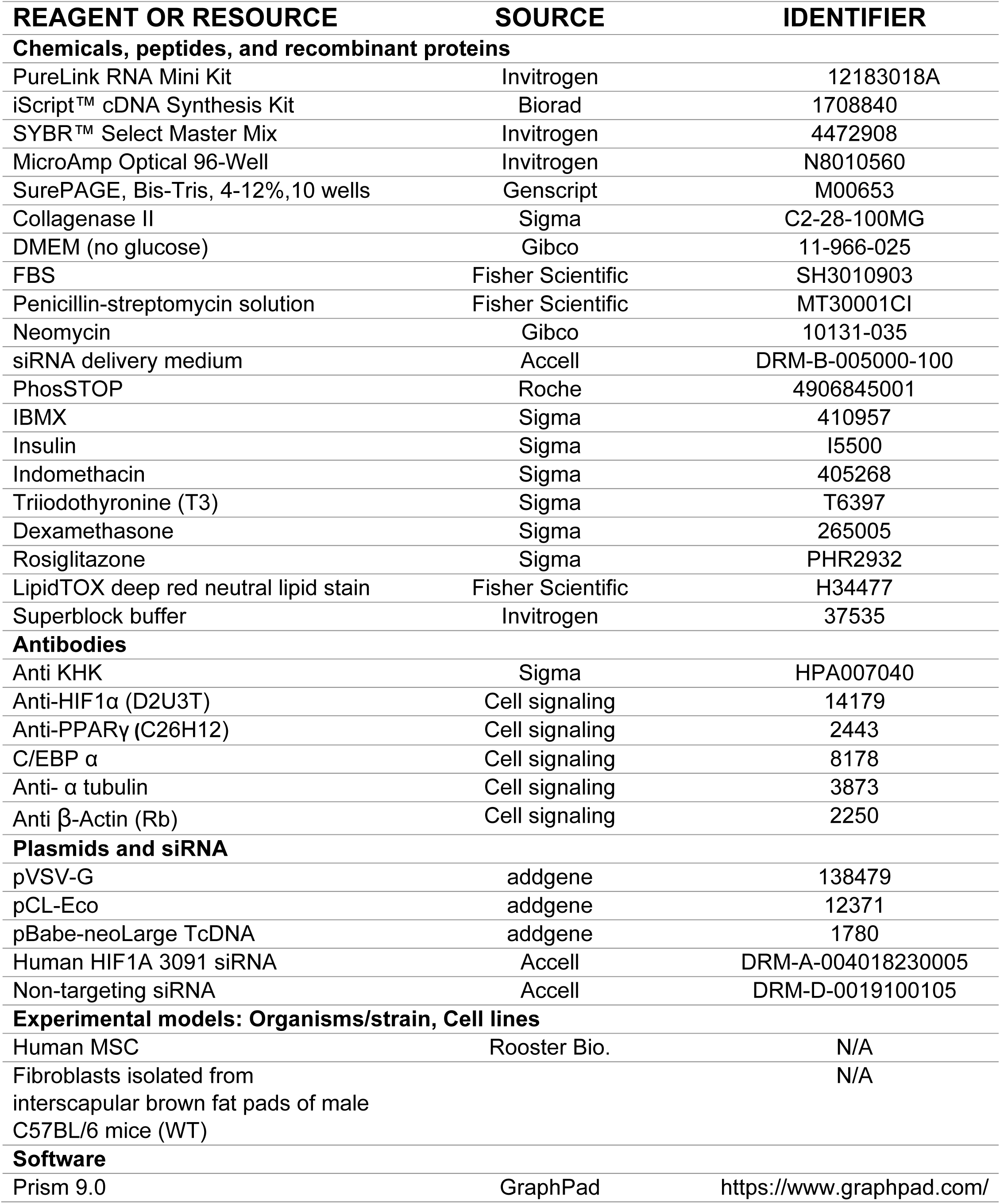
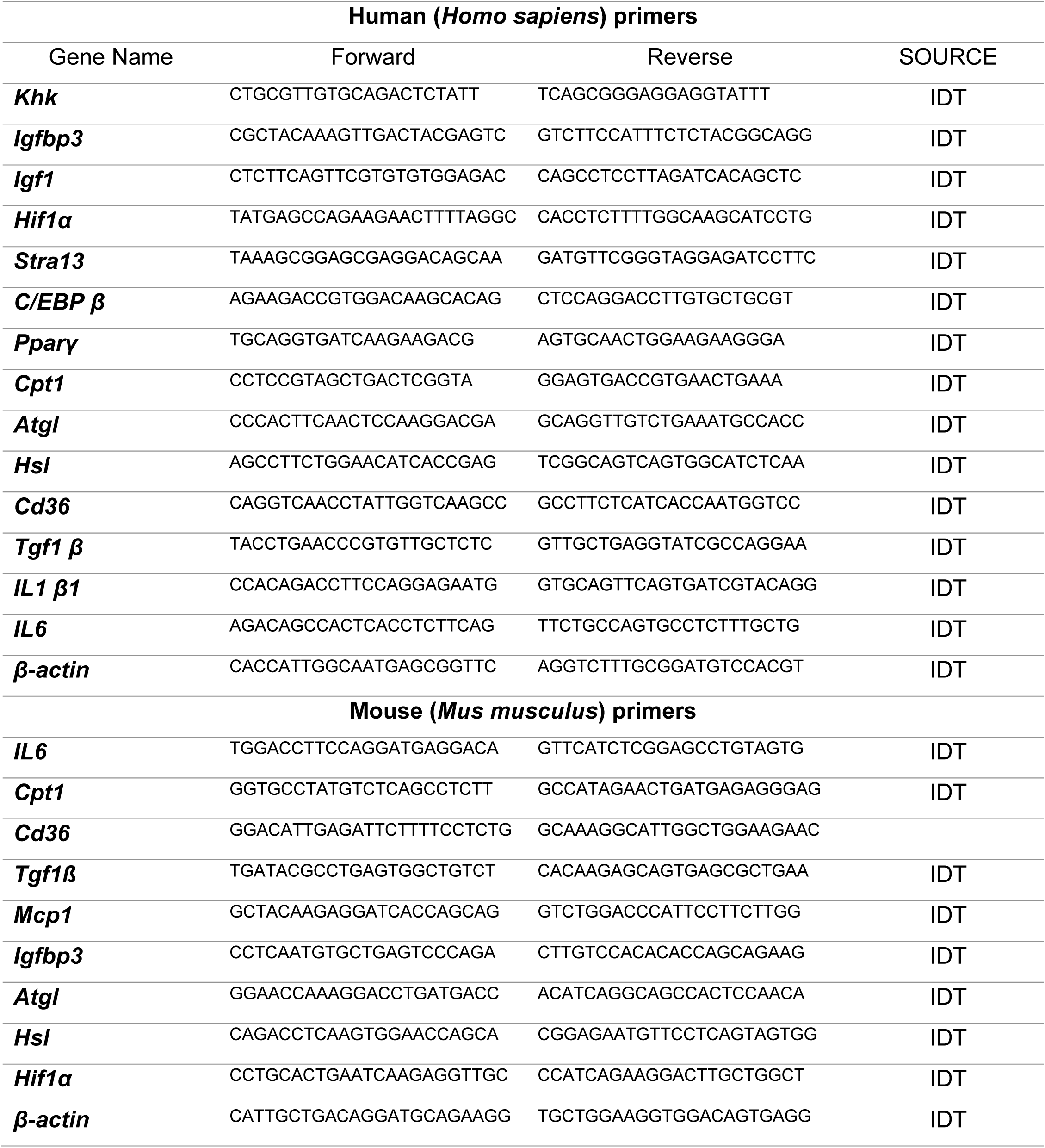
KEY RESOURCES TABLE.

## METHOD DETAILS

### MSC culture and differentiation

MSCs were cultured in Iscove’s modified Dulbecco’s medium (IMDM) supplemented with 10% fetal calf serum (FCS). The culture medium also contained 100 IU/ml of penicillin, 100 mg/ml of streptomycin (P-S) and 10 ng/ml of platelet-derived growth factor BB (PDGF-BB). Cells at 80-90% confluency were incubated for 3 weeks in adipogenic differentiation media which consisted of IMDM supplemented with 10% rabbit serum, 0.5 mM 3-isobutyl-1-methylxanthin (IBMX), 1 µM hydrocortisone, 0.1 mM indomethacin, and P–S. Media were changed every 3 days. MSCs were fixed with cold 10% formalin for 15 min, washed twice with PBS, and cytoplasmic triglyceride droplets were stained with BODIPY (Sigma) for 15 min at room temperature (RT). Cells were washed and mounted with Vectashield containing DAPI nuclear stain, table (1). Cells were observed under Leica fluorescent microscope (Leica DMi8 system).

### Isolation and immortalization of stromal vascular cells

Brown preadipocyte isolation and immortalization was previously described with modification^42–44^. Briefly, brown fat tissue was removed from interscapular region, minced into small piece, and digested in buffer including 1 mg/ml collagenase II at 37C for 30-40 min. digested tissues were then filtered through a 100μm cell strainer into a new 50ml sterile tube. Cells were then pelleted by centrifuge at 600 g for 5 min, and plated into a 6-well culture plate with DMEM/F12, 10% FBS, and 1% penicillin-streptomycin and cultured in 37C and 5% CO_2_ for overnight. Cells were washed with pre-warmed 1× PBS, then infected with retrovirus to express SV40 Large T antigen. The retroviral particles were packaged by 293T cells transfected with pVSV-G, pCL-Eco, and pBabe-neolarge TcDNA plasmids, table (1). After Infecting for 48 hours, cells were cultured in selection medium including with 450ug/ml neomycin. When reaching 80-90% confluence, cells were sub-cultured to a new 6-well plate. Cells were remained in neomycin selection for at least 7 days.

### Differentiation of mouse primary brown adipocyte

Immortalized preadipocytes were grown in the DMEM medium supplemented with 1g/L glucose, 10% FBS and 1% penicillin-streptomycin. When cells reached to 100% confluence, differentiation (day 0) was initiated with the induction medium with 1g/L glucose or 1g/L fructose supplemented with 0.5mM IBMX, 20 nM insulin, 125nM indomethacin, 1nM T3, 1μM dexamethasone, 1μM rosiglitazone for 48 hours. Cells were then incubated in the maintenance medium with 1g/L glucose or 1g/L fructose supplemented with 20nM insulin, 1nM T3 for 12 days.

Live cell lipid droplet staining was achieved by HCS LipidTOX deep red neutral lipid stain and imaged with fluorescence microscopy (Leica DMi8 system).

Differentiated mouse adipocytes were cultured in Dulbecco’s modified Eagle’s medium (25mmol/l glucose), supplemented with 10% (vol/vol) FBS, 1% penicillin, and streptomycin, and 1% Gultamax. Medium was exchanged every 3 days, and cells were trypsinized and reseeded at 1:4 dilution when 80-90% confluence was reached, all reagent designated in table (1).

### Cytokines measurement in the conditioned medium

Conditioned media were collected from either human MSCs or from mouse fibroblasts (pre-adipocytes). Samples were assayed for cytokines by the University of Maryland, Baltimore Cytokine Core Lab according to manufacturer’s directions.

Briefly, a 96 well plate (Greiner) was wet with 200ul of Assay Buffer and placed on a shaker for 10 minutes. The plate is then decanted and 25ul of Assay Buffer or appropriate buffer is added to each well and 25ul of standard/sample/control was added to the appropriate wells. Samples were run in duplicate for all samples. Then 25ul of a mixture containing required cytokines (1:50 dilution) that have been conjugated to beads is added. All plates contain at minimum high and low control in order to determine the validity of the plates. The plate is then placed on a shaker, at 4C overnight. The plate was then placed on a magnetic washer, 200ul of Wash Buffer added to each well, the plate was set on a shaker at 500 rpm for 1 minute, and repeated an additional two times. After the last decanting step 25ul of detection antibody is added and the plate is placed on a shaker for one hour at room temp. Then 25ul of Phycoerythrin (1:25 dilution) is added to each well and the plate is placed back on the shaker for 30 minutes. The plate is then washed three times and 150ul of Sheath Fluid is added to each well. The plate is then read using a Luminex MagPix reader. The data is then calculated using Luminex’s exponent Software.

### RNA extraction and qPCR

Total RNA was extracted from 2×10^6^ HepG2 or Huh7 cells using PureLink RNA Mini Kit (Invitrogen) according to manufacturer instructions. iScript™ cDNA Synthesis Kit (Biorad) was used to convert one microgram of high purity RNA into cDNA in a total reaction volume of 20 µL. The rection volume was diluted to 200 uL to reach (5 ng/ µL) .2 µL was used for qPCR (10 ng) per reaction.

Two microliter (2 µL ∼10 ng of cDNA) of this cDNA was used for subsequent PCR amplification with SYBR™ Select Master Mix (cat. 4472908, Invitrogen) in QuantStudio 5 (ThermoFisher scientific) using the following specific primers for human, Table (2) or for mice, Table (3) were purchased from Integrated DNA Technology, IDT.

### Western blot analysis

Cells were lysed in RIPA buffer, 150 mM NaCl,50 mM HEPES, pH 7.6, containing 0.1% of (100× Protease inhibitor) and one tablet of PhosSTOP. Equal amounts of protein samples were blocked in 4X loading buffer and run in 4-12% SurePAGE™, Bis-Tris gel then transferred to a PVDF membrane. After blocking with superblock buffer, Immunoprobing the membrane with specific antibodies designated in the table (1).

### Statistical analysis

Data were analyzed with GraphPad Prism 9 software (GraphPad Software, USA, table1) and presented as Mean ±SEM. We used one-way ANOVA followed by Post HOC Tukey test for multiple comparison. Statistical significance is indicated as figures using the following denotations, \**P*<0.05, \*\**P*<0.01, \*\*\**P*<0.001, \*\*\*\**P*<0.0001.

## Results

### Cellular growth was modulated by fructose

Standard IMEM growth medium includes 25 mM glucose supplemented with 10 ng/L of PDGF and 10% FCS. We investigated the cellular growth dynamics under increasing concentrations of either glucose or fructose. Growth dynamics were dampened when MSCs grown in fructose compared to glucose at equivalent concentrations, except for 25 mM fructose (Fig. 1, A-B).

**Fig. 1.**
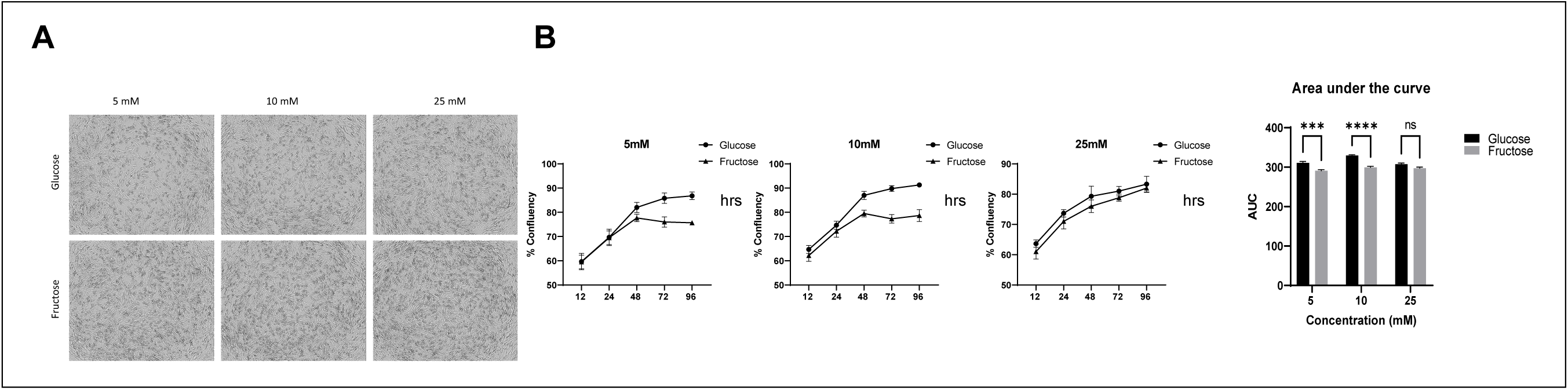
Fructose decreases cellular growth. The growth curves of MSCs at concentrations of 5, 10, and 25 mM in either glucose or fructose, (A &B). MSCs grow faster in glucose when compared to fructose at equivalent concentrations.

### Cytokine production changes in fructose compared to glucose

The striking differences in growth dynamics between fructose and glucose prompted us to screen out the cytokines released by MSCs in the conditioned medium. Those cytokines influence either cell proliferation or senescence. Interestingly, MSCs under fructose significantly release more senescence markers such as IL4, IL8, and IL10^21–24^. Notably, the increase of GM-CSF is associated with the differentiation into monocyte lineage^25^. Conversely, GDF-15 secretion was reduced indicating the reduction in cellular growth^26^ (Fig.2-A). This shift in cytokine secretion highlights the impact of fructose on MSC, leading to alterations in proliferation.

**Fig. 2.**
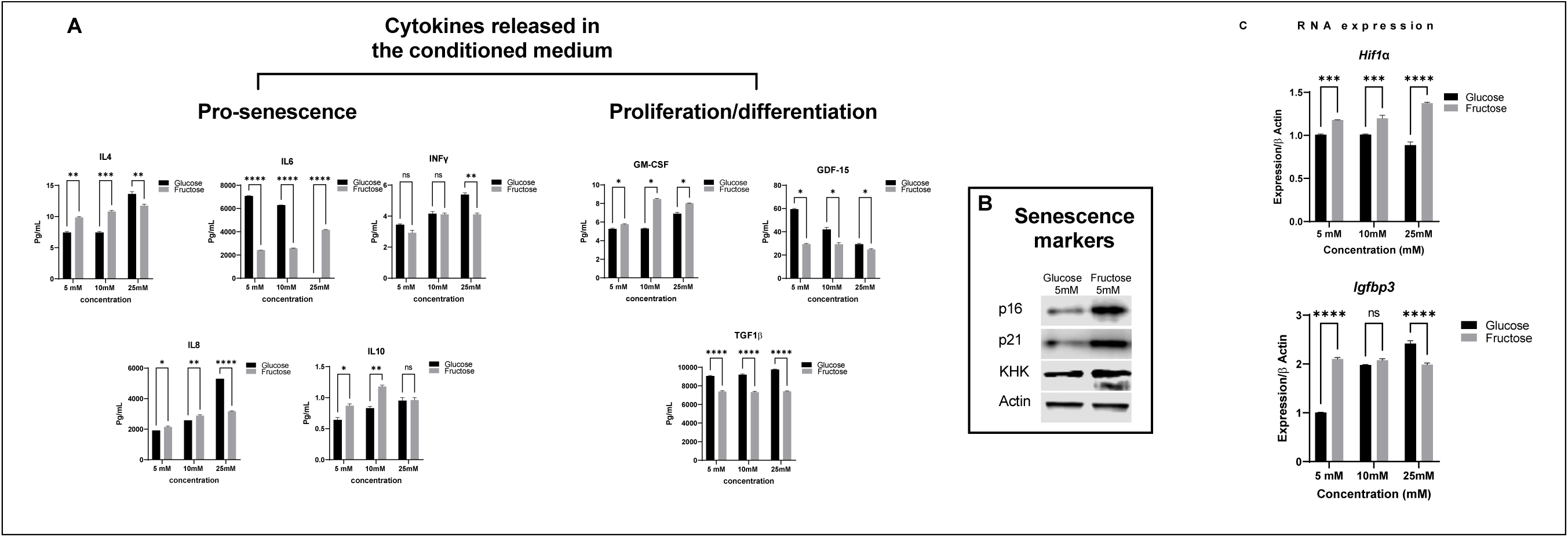
Fructose modulates cytokine production. Eight different cytokines were measured in the conditioned medium. These cytokines are either activators or suppressors for cell proliferation (A). Western blot analysis of senescence marker, P16 and P21 as well as KHK were elevated (B). *Hif1α* is a marker for cellular stress. Its expression increased in the fructose treated MSCs in an oxygen independent mechanism. This was associated with the increase of Igfbp3 expression (C).

Consequently, we asked if fructose modulates the expression of cell senescence markers, specifically, P16 and P21. P16INK4a or P16, is an inhibitor of cyclin-dependent kinases and plays a critical role in cell cycle regulation, enforcing the G1 phase cell cycle arrest in senescent cells^27–29^. Similarly, P21CIP1/WAF1 or P21 prevents cell cycle progression^29,30^. Western blot analysis of MSCs cultured in 5mM fructose showed a significant increase in immunosignal of both markers (Fig.2-B). Consequently, fructose not only enhances the production of senescence-associated cytokines but also activates cell senescence markers in MSCs.

### Cellular stress caused by fructose upregulates *Hif1α* expression

Changes in the microenvironment can exert a profound impact on cell vitality. For instance, high levels of glucose or fructose^32^ as well as pollutants have the potential to induce stress factors like Hif1α, even under normoxic conditions, ultimately leading to senescence^31,32^. Notably, one of the significant challenges in stem cell transplantation is cell death induced by Hif1α activation^33^. Accordingly, we investigated transcript levels of *Igfbp3* as a direct indicator of *Hif1α* activation^34,35^, as a consequence of fructose as the sole carbon source. We found that *Hif1α* was upregulated in the fructose treated cells. Additionally, *Igfbp3* mRNA levels were upregulated at 5mM and 25mM fructose (Fig. 2-C). Therefore, *Igfbp3* and *Hif1α* expression were modulated when MSCs grown under fructose.

### Oxygen tension and sugar supplementation boosted cytokine expression in MSCs

We showed that substituting glucose with fructose imposes metabolic stress on the growth of stem cells and alters the production of cytokines. This stress induces the expression of *Hif1α* and *Igfbp3* regardless of the presence of oxygen. Notably, stem cells can be found in tissues that receive low levels of oxygen such as bone marrow stem cells ^1,36^ and stem cells derived from expanding white adipose tissue^37,38^. This prompted us to further investigate the response of MSCs under different oxygen tensions, specifically comparing normoxia (20% O_2_) and hypoxia (1% O_2_).

Surprisingly, exposure to hypoxia did not result in a significant change in the transcription levels of *Hif1α* itself (Fig. 3-A). However, at the protein level, there was a comparable increase in HIF1α, that was observed in fructose-treated MSCs, irrespective of the oxygen tension(Fig. 3-B). Notably, the combination of fructose and hypoxia had a substantial effect on the upregulation of HIF1α, *Igfbp3*, *Igf1*, and *Khk*, a key enzyme in fructose metabolism (Fig.3-A). Interestingly, the measurement of IGFBP3 and IGF1 released by MSCs in the culture medium closely matched the mRNA expression levels. (Fig. 3-C). These results provide compelling evidence that establishes a direct relationship between *Hif1α* and *Igfbp3* expression during hypoxic conditions in MSCs.

**Fig. 3.**
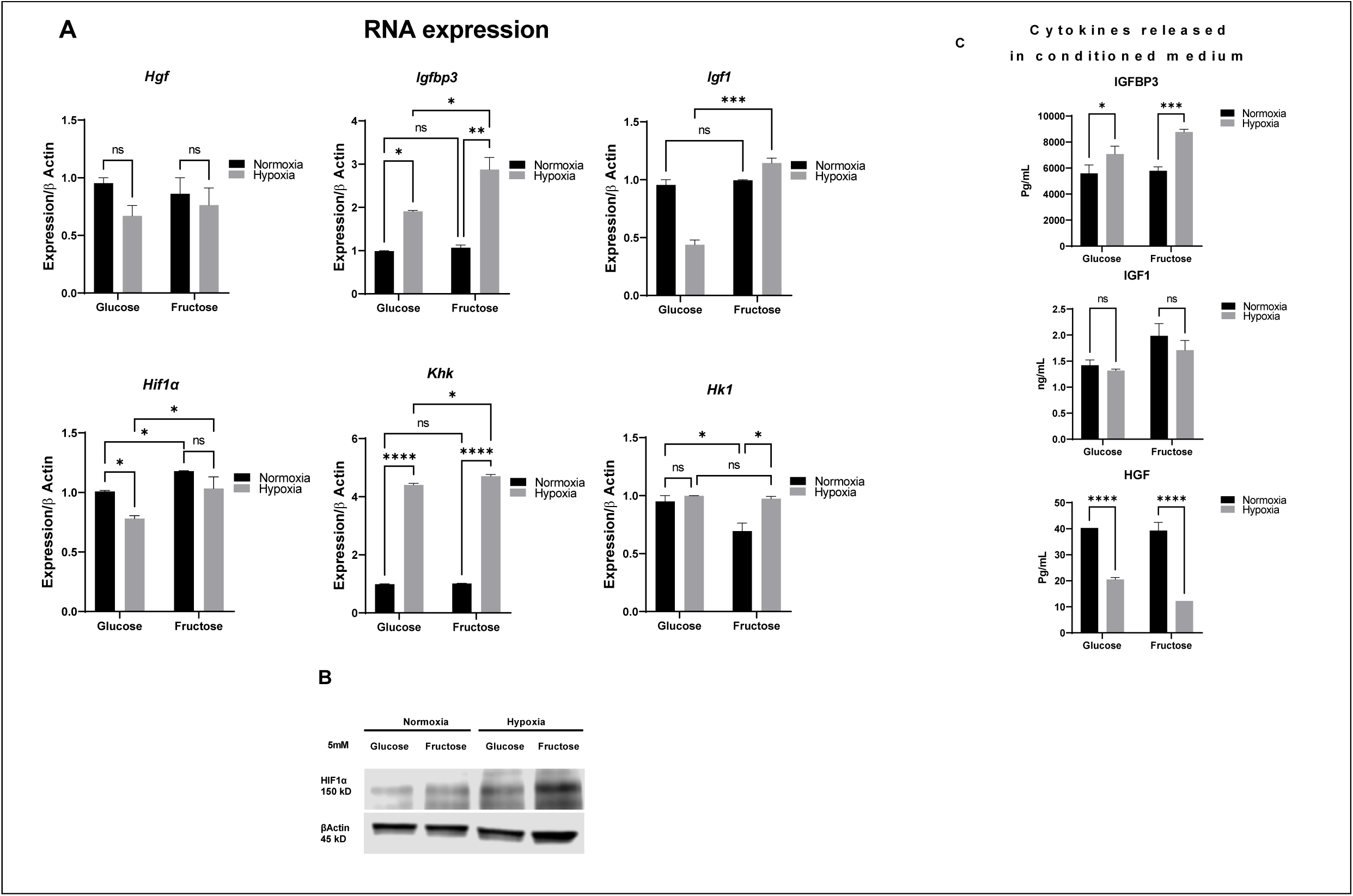
Low oxygen and sugar supplement changes cytokine expression in MSCs. We incubated MSCs in either 5mM of glucose or fructose under two oxygen levels, 20% and 1% to induce hypoxia. While the mRNA expression of *Hif1α* did not change (A), the immunosignal of HIF1a protein was induced in fructose treated MSCs (B). *Igfpb3*, *Igf1*, and *Khk* were significantly upregulated in the presence of fructose and hypoxia (A). Interestingly, the increase in concentrations of IGFBP3 and IGF1 in the medium closely mirrored the observed changes in expression levels (C).

### Knocking down *Hif1α* reduced *Igfbp3* expression in MSC grown in either glucose or fructose

Our study revealed that the interplay between *Hif1α* and *Igfbp3* is influenced by both fructose and oxygen levels. This prompted us to delve deeper by knocking down *Hif1α* using small interfering RNA (siRNA). The knockdown was associated with reduction of *Igfbp3* expression (Fig. 4-A) and in the culture medium but not in fructose (Fig. 4-B). Interestingly, the levels of IGF1 released into the culture medium remained unchanged (Fig. 4-B). These results offer compelling confirmation that *Hif1α* functions as an upstream regulator for *Igfbp3* expression in MSCs.

**Fig. 4.**
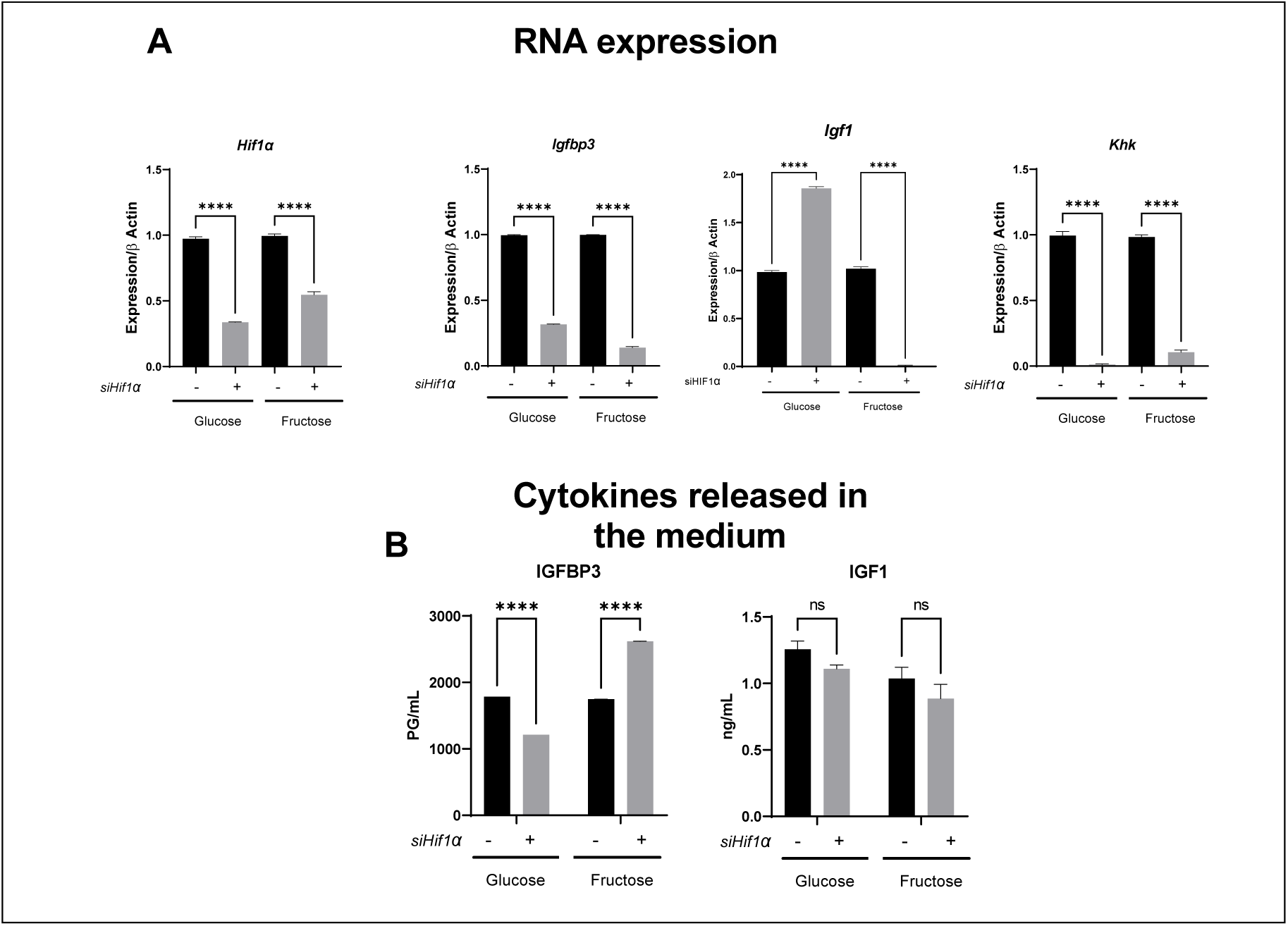
knocking down *Hif1α* reduced *Igfbp3* expression and cytokines released in the medium. *siHif1α* significantly reduced the expression of *Hif1α, Khk,* and *Igfbp3* (A). IGFBP3 measured in the glucose containing medium was reduced but not in fructose. Additionally, there was no observed change in the levels of IGF1.

### Substituting glucose with fructose does not promote adipogenesis

The profound stress that fructose exerts on MSCs proliferation and cytokine production motivated us to further explore the impact of fructose on differentiation. To examine the impact of fructose on adipogenesis, we employed two distinct cell types: human multipotent mesenchymal stem cells (MSCs) and primary fibroblasts isolated from brown fat tissue in mice^42–44^. Standard induction medium contains 25 mM glucose. Hence, we investigated the effect of substituting glucose with fructose in differentiation medium at various concentrations (5, 10, and 25 mM).

In glucose, significant MSC differentiation was observed, visualized as accumulation of lipid droplets by BODIPY staining (Fig. 5-A). Conversely, when MSCs were exposed to fructose, adipocyte differentiation and the subsequent accumulation of lipids displayed a noticeable reduction, even though cell viability remained largely unaffected.

**Fig. 5.**
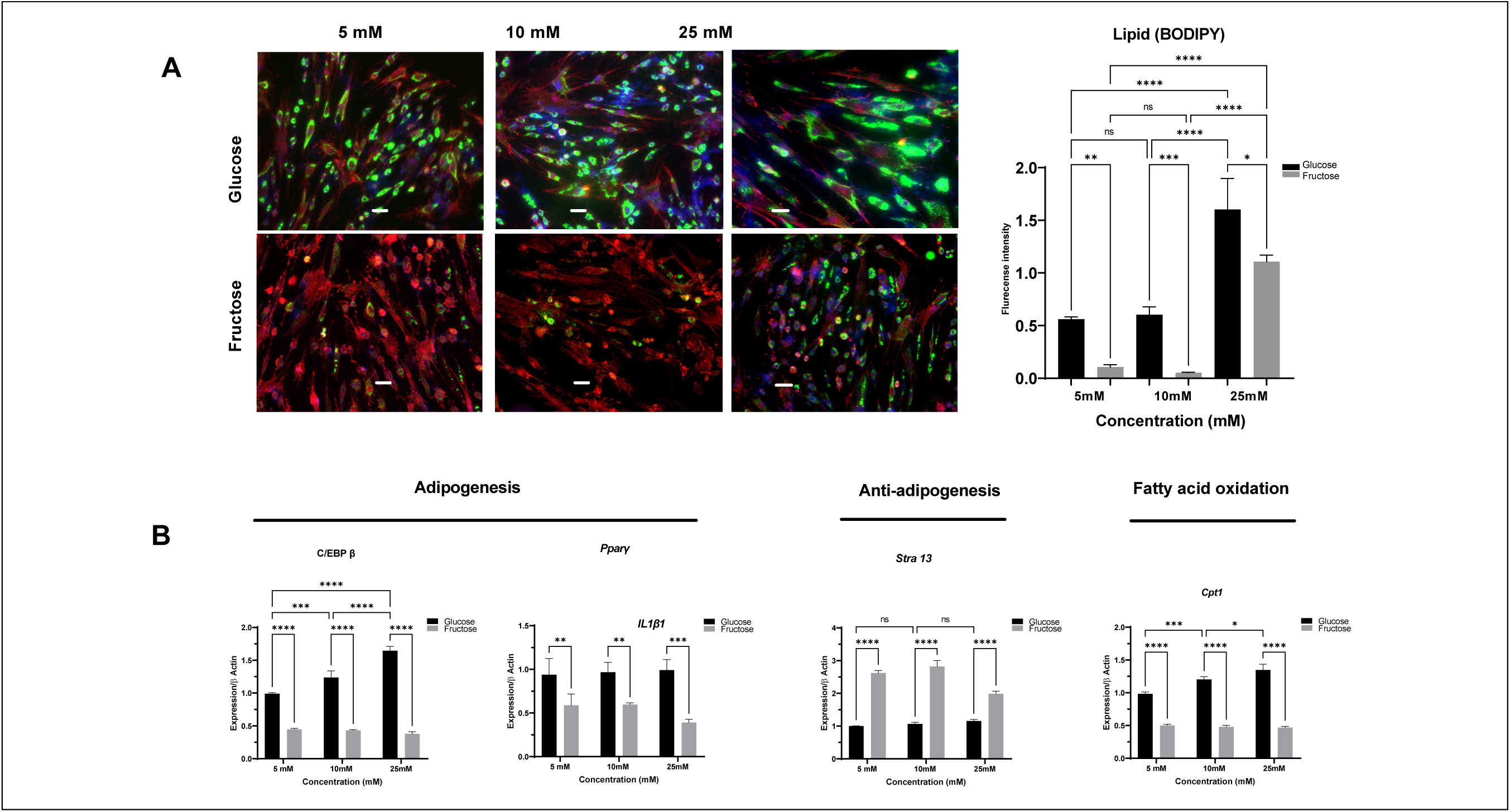
Fructose modulated Adipogenesis in MSCs Compared to Glucose. (A) MSCs were cultured in adipogenic differentiation medium supplemented with either glucose or fructose. After 14 days of differentiation, cells were fixed with 4% PFA, and lipid accumulation was visualized using BODIPY stain (green), F-Actin (red), and nuclei (blue). A) Lipid accumulation was observed across a spectrum of glucose concentrations, ranging from 5 mM to 25 mM. In contrast, the induction of adipogenic differentiation was only achieved at the 25 mM fructose concentration (B). Fructose treated cells showed elevated levels of IL1β1 and a reduction in Cpt1 expression.

Notably, MSCs exposed to fructose showed a significant decrease in the expression levels of adipogenic markers, specifically *C/EBPβ* and *Pparγ* as well as *Hif1α*. In contrast, *Stra13*, a transcription factor often associated with the repression of adipogenesis was upregulated. Intriguingly, fructose also increased the expression of IL1β1, and a decrease in Cpt1, indicating a reduced capacity for fatty acid oxidation. (Fig. 5-B). Taken together, fructose failed to promote adipogenic differentiation of MSCs but, rather, enhanced the expression of proinflammatory cytokines.

### Mouse primary adipocytes failed to differentiate under fructose

Beyond MSC, fructose also impaired adipogenic differentiation of mouse pre-adipocytes. At day zero of differentiation, cells secrete higher levels of senescence associated cytokines, specially, IL1α1, IL6, IL8, MCP1, and TNF1 α (Fig. 6-A). Notably, these cytokines exert a substantial inhibitory effect on adipogenesis after twelve days of differentiation. This was clearly shown by the remarkable reduction in lipid accumulation in the fructose-treated cells when compared to glucose (Fig. 6-B). This reduction in lipid accumulation coincided with a decrease in PPARγ, a key transcription factor for adipogenesis. The immunosignal of HIF1α was also reduced in the fructose-treated cells, while KHK was upregulated (Fig. 6-C). Interestingly, fructose induced lipolysis via upregulation of hormone sensitive lipase (Hsl) (Fig. 6-E). This was associated with upregulation of two cytokines, IL6 and Tgf1β (Fig. 6-D), both recognized for their capacity to stimulate lipolysis. Surprisingly, Cpt1 expression was upregulated indicating an enhancement of fatty acids oxidation (Fig.6-F). In sum, fructose failed to support differentiation of murine preadipocytes, but instead induced lipolytic activity through the expression of IL6 and Tgf1β cytokines, along with elevated fatty acid oxidation.

**Fig. 6.**
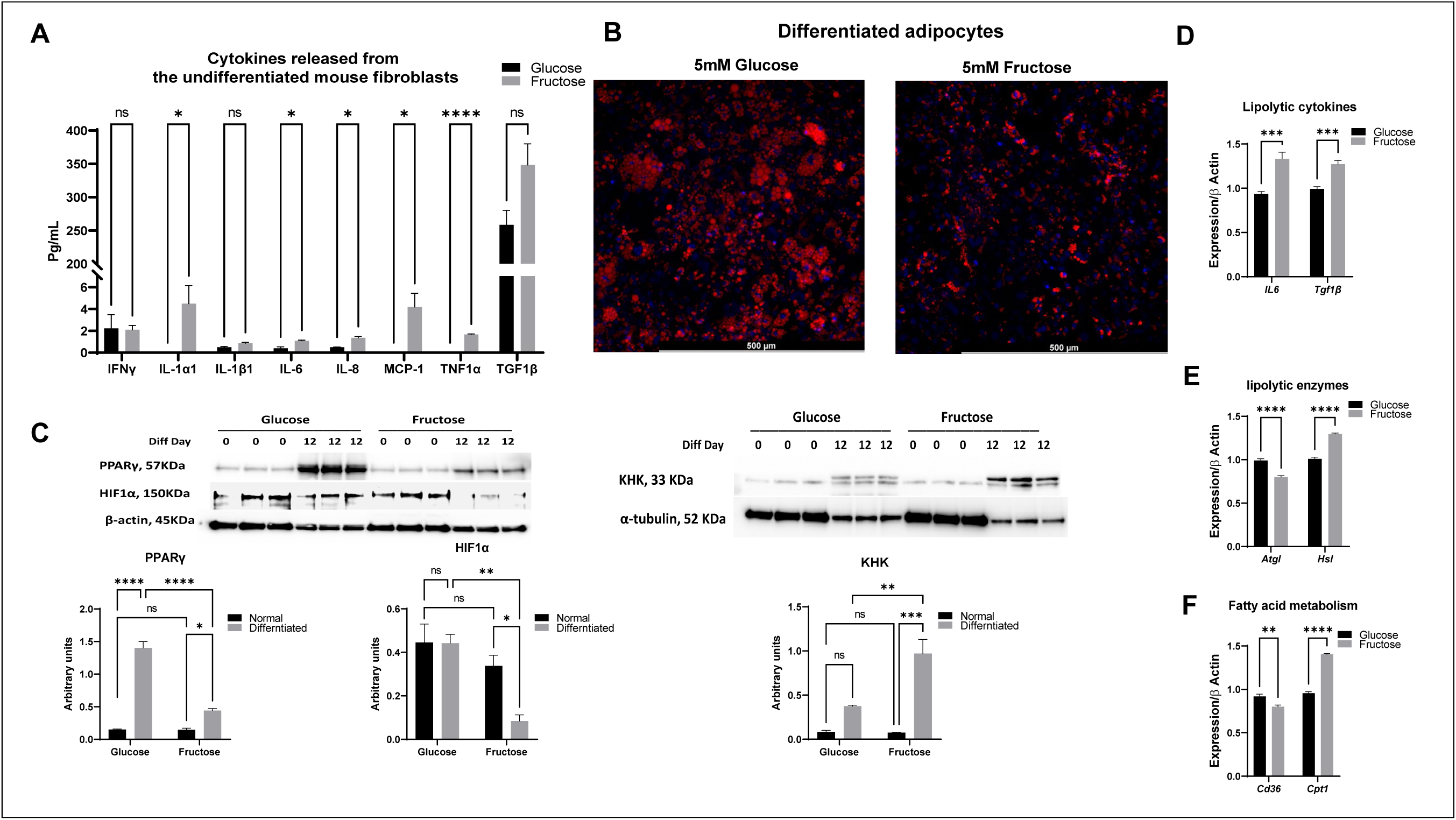
Fructose hindered adipogenesis in primary pre-adipocytes. Primary mouse fibroblasts showed elevation of senescence associated cytokines when exposed to fructose (A). The primary fat cells were cultured in adipogenic differentiation medium supplemented with either 5 mM glucose or fructose. After 12 days of differentiation, cells were fixed with 4% PFA, and lipid accumulation was visualized using neutral red stain red, and nuclei, blue (B). Western blot showing PPARγ, a major adipogenic factor at day 0 and at day 12, and HIF1α. Both proteins were downregulated in the fructose-treated cells (C). An Increase in Hsl, a lipolytic enzyme (E) was associated with upregulation in IL6 and TGF1β expression (D) concomitant with Cpt1 upregulation, a maker for FA oxidation (F).

## Discussion

Our bodies contain various types of stem cells that play a vital role in tissue regeneration and wound healing^45,46^. They greatly vary not only in their origins but also in their potency^6^. This can be attributed partially to the special microenvironment surrounding them that contains the required nutrient, oxygen level, and cytokines suitable for survival and differentiation^4,6^. Any change in the pericellular environment will consequently modulate cell signals, cytokines, that will act directly on the cell itself (autocrine effect) or neighboring cells (paracrine effect)^3,4,6^. Therefore, preserving the pericellular environment of stem cells is crucial for their normal function.

For instance, elevated glucose and fructose levels in diabetic patients exert significant stress on stem cells within various organs^47,48^. High fructose can generate toxic metabolites such as uric acid that in turn elevates ROS production^49^. Additionally, fructose metabolism leads to ATP depletion that triggers apoptosis in primary hepatocytes^50,51^. This stress leads to increased expression of Hif1α, a factor known for promoting adverse effects in multiple organs, including angiogenesis in retina^52^ and white adipose tissue^53^, and osteoporosis^1,36^. HIF-1α is subject to rapid ubiquitination under normal oxygen conditions^54^. However, in hypoxic environments or when exposed to prolyl hydroxylases (PHDs) inhibitors such as ROS, HIF-1α becomes more stable and translocate into the nucleus^54^. Hence, elevated hexose and ROS production can induce stress leading to HIF1a stabilization and modulation of stem cells viability.

Surprisingly, we observed that replacing glucose with fructose also subjects stem cells to stress, resulting in increased Hif1a expression and stability. This led to a reduction in cell proliferation, and alterations in cytokine production. The increase of Hif1a is evident through its induction of Igfbp3 expression and secretion^34^. Notably, Igfbp3 triggers apoptotic pathway and inhibits cellular proliferation^35,55^. Conversely, we found that Igfbp3 expression decreased when Hif1a was knocked down, regardless of the sugar type used. The induction of Hif1a influence the production of cytokines associated with the senescence-related phenotype^33,56^, aligning with our findings that demonstrate an increase in P16 and P21 immunosignals. Interestingly, we showed that IL4, IL8, and IL10 were increased in the fructose containing medium along with the reduction in GDF15, a growth factor that supports proliferation^26^. These three interleukins are associated with senescence phenotype^21,24^. Hence, fructose exerts stress on stem cells associated with increased Hif1a and activation of senescence associated phenotype.

The previous findings prompted us to ask a question whether this phenomenon extends to differentiation. To address this, we conducted a comparative study of the adipogenic differentiation potential of human MSCs, which are multipotent, and mouse fibroblasts isolated from the interscapular brown adipose tissue. These mouse fibroblasts are more specialized and can only differentiate into brown adipocytes^57^. This contrast in their differentiation capabilities offers an excellent model for examining how the replacement of glucose with fructose might influence their differentiation.

Interestingly, fructose failed to induce differentiation of human MSCs as well as mouse fibroblasts into mature adipocytes compared to glucose. Key markers of adipogenesis, including C/EBPβ, and PPARγ^37^ were upregulated in fructose-treated MSCs but did not reach the threshold required for adipogenesis when compared to glucose (supplementary fig. 1-A). Conversely, we showed that fructose induces undifferentiated mouse fibroblasts to release cytokines associated with senescence, including IL1α1, IL6, IL8, MCP1, and TNF1α. This observation prompted us to investigate further by differentiating these cells into adipocytes. Intriguingly, in fructose-treated mouse fibroblasts, there was a remarkable increase in IL6, a well-known cytokine that stimulates lipolysis^58–60^. This was supported by the significant upregulation of *Hsl* concomitant with the increase in *Cpt1* expression. These changes suggest that these cells were undergoing lipolysis, with fatty acids being oxidized to generate energy.

Surprisingly, human MSCs exhibited elevated IL1β1 and IL6 expression also known for inducing lipolysis^57,61^. Notably, the expression of lipolytic enzymes Atgl and Hsl remained unchanged while Cpt1 was downregulated. Hence, fructose impedes adipogenic differentiation by elevating the expression of cytokines that promote both lipolysis and the oxidation of fatty acids.

In conclusion, our study has revealed that altering the cultural conditions through changes in hexose levels and oxygen tension places considerable stress on stem cells. This stress not only leads to shifts in cytokine composition but also activates pathways associated with senescence or inhibits the maturation of stem cells into adipocytes. This study opens the door to further investigations into the mechanisms governing stem cell response to their microenvironments.

---

**Supplemental Figures 1.**
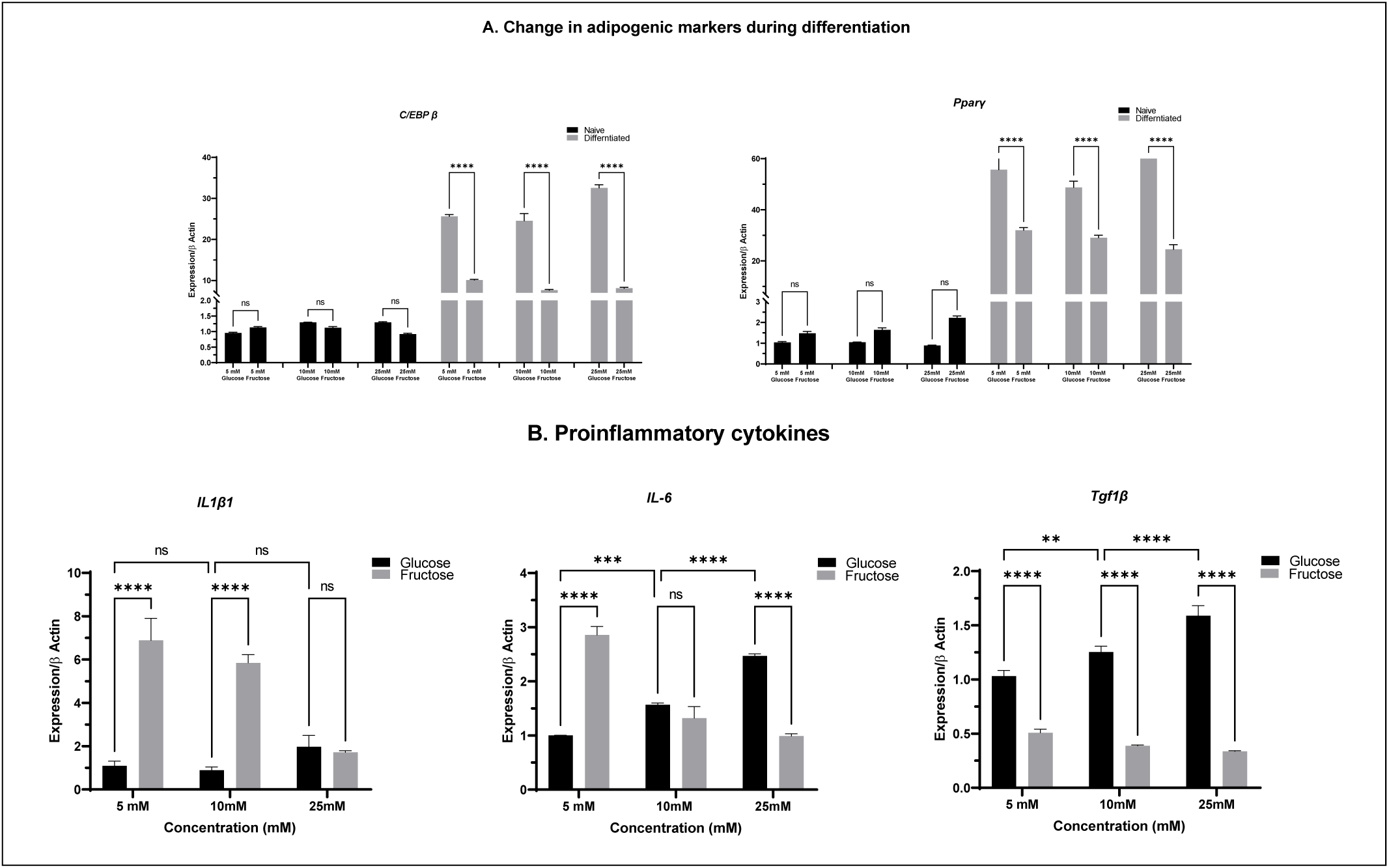
Fructose does not support MSCs to differentiate into mature adipocytes.

**Supplementary fig (1).**
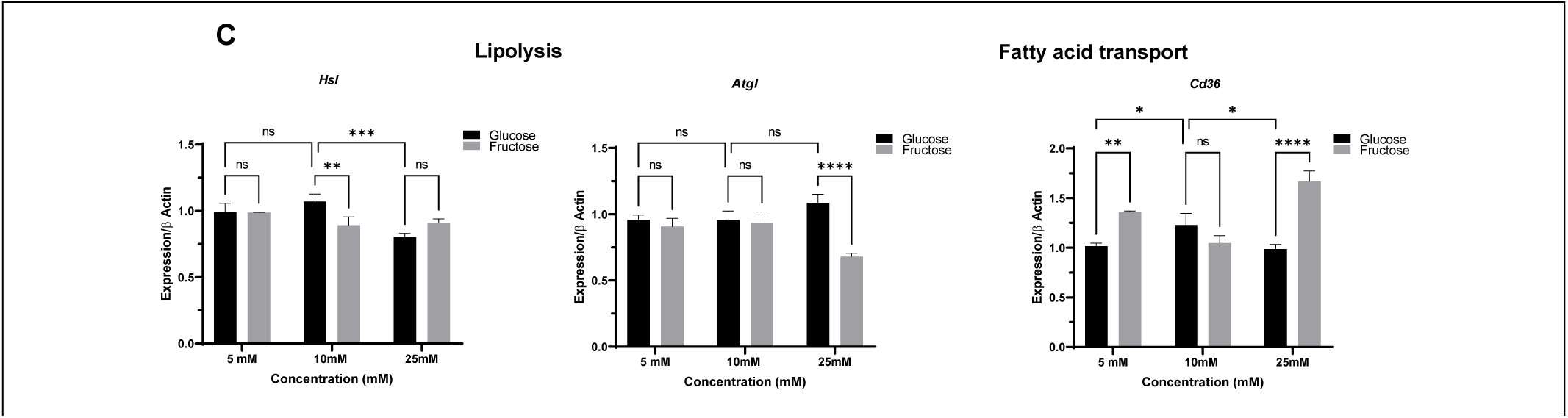
MSCs under fructose showed significant upregulation of C/EBPβ and Pparγ but not enough to induce adipogenesis (A). Proinflammatory cytokines IL1β1 and IL6 are known for their ability to induce lipolysis as well as Cd36, a fatty acid transporter, showed significant upregulation in MSCs differentiated in fructose (B). Conversely, Hsl, a lipase that is directly activated via interleukin as well as Atgl, remained unchanged (C).

## METHOD DETAILS

### MSC culture and differentiation

MSCs were cultured in Iscove’s modified Dulbecco’s medium (IMDM) supplemented with 10% fetal calf serum (FCS). The culture medium also contained 100 IU/ml of penicillin, 100 mg/ml of streptomycin (P-S) and 10 ng/ml of platelet-derived growth factor BB (PDGF-BB). Cells at 80-90% confluency were incubated for 3 weeks in adipogenic differentiation media which consisted of IMDM supplemented with 10% rabbit serum, 0.5 mM 3-isobutyl-1-methylxanthin (IBMX), 1 µM hydrocortisone, 0.1 mM indomethacin, and P–S. Media were changed every 3 days. MSCs were fixed with cold 10% formalin for 15 min, washed twice with PBS, and cytoplasmic triglyceride droplets were stained with BODIPY (Sigma) for 15 min at room temperature (RT). Cells were washed and mounted with Vectashield containing DAPI nuclear stain. Cells were observed under Leica fluorescent microscope (Leica DMi8 system).

### Isolation and immortalization of stromal vascular cells

Brown preadipocyte isolation and immortalization was previously described with modification^42–44^. Briefly, brown fat tissue was removed from interscapular region, minced into small piece, and digested in buffer including 1 mg/ml collagenase II at 37C for 30-40 min. digested tissues were then filtered through a 100μm cell strainer into a new 50ml sterile tube. Cells were then pelleted by centrifuge at 600 g for 5 min, and plated into a 6-well culture plate with DMEM/F12, 10% FBS, and 1% penicillin-streptomycin and cultured in 37C and 5% CO_2_ for overnight. Cells were washed with pre-warmed 1× PBS, then infected with retrovirus to express SV40 Large T antigen. The retroviral particles were packaged by 293T cells transfected with pVSV-G, pCL-Eco, and pBabe-neolarge TcDNA plasmids. After Infecting for 48 hours, cells were cultured in selection medium including with 450ug/ml neomycin. When reaching 80-90% confluence, cells were sub-cultured to a new 6-well plate. Cells were remained in neomycin selection for at least 7 days.

### Differentiation of mouse primary brown adipocyte

Differentiated mouse adipocytes were cultured in Dulbecco’s modified Eagle’s medium (25mmol/l glucose), supplemented with 10% (vol/vol) FBS, 1% penicillin, and streptomycin, and 1% Gultamax. Medium was exchanged every 3 days, and cells were trypsinized and reseeded at 1:4 dilution when 80-90% confluence was reached.

### RNA extraction and qPCR

Two microliter (2 µL ∼10 ng of cDNA) of this cDNA was used for subsequent PCR amplification with SYBR™ Select Master Mix (cat. 4472908, Invitrogen) in QuantStudio 5 (ThermoFisher scientific) using the following specific primers for human, Table (1) or for mice, Table (2) were purchased from Integrated DNA Technology, IDT.

### Western blot analysis

Cells were lysed in RIPA buffer, 150 mM NaCl,50 mM HEPES, pH 7.6, containing 0.1% of (100× Protease inhibitor) and one tablet of PhosSTOP (No. 4906845001). Equal amounts of protein samples were blocked in 4X loading buffer and run in 4-12% SurePAGE™, Bis-Tris gel (No. M00653) then transferred to a PVDF membrane. After blocking with superblock buffer (INV-37535), Immunoprobing the membrane with specific antibodies designated in the table.

### Statistical analysis

Data were analyzed with GraphPad Prism 9 software (GraphPad Software, USA) and presented as Mean ±SEM. We used one-way ANOVA followed by Post HOC Tukey test for multiple comparison. Statistical significance is indicated as figures using the following denotations, \**P*<0.05, \*\**P*<0.01, \*\*\**P*<0.001, \*\*\*\**P*<0.0001.

